# Sensitivity analysis identifies the factors driving the enzymatic saccharification of lignocellulose

**DOI:** 10.1101/2023.09.22.559023

**Authors:** Partho Sakha De, Jasmin Theilmann, Adélaïde Raguin

## Abstract

Corn stover is the most abundant form of crop residue that can serve as a source of lignocellulosic biomass in biorefinery approaches, for instance for the production of bioethanol. In such biorefinery processes, the constituent polysaccharide biopolymers are typically broken down into simple monomeric sugars by enzymatic saccharification, for further downstream fermentation into bioethanol. However, the recalcitrance of this material to enzymatic saccharification invokes the need for innovative pre-treatment methods to increase sugar conversion yield. Here, we focus on experimental glucose conversion time-courses for corn stover lignocellulose that has been pre-treated with different acid-catalysed processes and intensities. We identify the key parameters that determine enzymatic saccharification dynamics by performing a Sobol’s sensitivity analysis on the comparison between the simulation results from our complex stochastic biophysical model, and the experimental data that we accurately reproduce. We find that the parameters relating to cellulose crystallinity and those associated with the cellobiohydrolase activity are predominantly driving the enzymatic saccharification dynamics. We confirm our computational results using mathematical calculations for a purely cellulosic substrate. On the one hand, having identified that only five parameters drastically influence the saccharification dynamics allows us to reduce the dimensionality of the parameter space (from nineteen to five parameters), which we expect will significantly speed up our fitting algorithm for comparison of experimental and simulated saccharification time-courses. On the other hand, these parameters directly highlight key targets for experimental endeavours in the optimisation of pre-treatment and saccharification conditions. Finally, this systematic and two-fold theoretical study, based on both mathematical and computational approaches, provides experimentalists with key insights that will support them in rationalising their complex experimental results.

**SIGNIFICANCE**
The recalcitrance of lignocellulosic biomass to enzymatic saccharification processes necessitates innovative biorefinery pre-treatment processes with the goal of increasing glucose conversion yields. We implement a systematic Sobol’s sensitivity analysis on the comparison between the simulation results from our complex stochastic biophysical model, and experimental glucose conversion time-courses data for pre-treated corn stover lignocellulose that we accurately reproduce with our model. Our results, reinforced with analytical calculations, highlight the key parameters which govern the dynamics of enzymatic saccharification of lignocellulose. Thus, this study does not only drastically reduce the model parameter space from nineteen free parameters to only five key ones, but it also provides valuable insights that can guide future experiment design efforts, and the rationalisation of complex experimental datasets.

## INTRODUCTION

Global challenges of energy shortage demand a major transition in our energy landscape from being predominantly fossil fuel based to more renewable sources. In addition to hydroelectricity, wind energy, tidal energy, solar energy, and geothermal energy, biofuels produced from cellulosic and lignocellulosic materials, such as agricultural and forestry residues, energy crops, or aquatic biomass, are major candidates for the replacement of fossil fuel energy sources, especially in the transport sector (1). Globally, agricultural systems generate over 200 billion tonnes of crop residues per year, most of which ends up being used as animal feed or being simply disposed of, hence wasting its significant potential as a valuable bioresource (2, 3). The most common leftover parts include corn stover, rice straw, wheat straw, and sugarcane bagasse, with corn stover being the most abundant one (4). This biomass is mainly composed of lignocellulose, consisting of 30-44% cellulose, 30-50% hemicellulose, and 8-21% lignin, depending on its source (3, 5). For the production of biofuels, the constitutive cellulose biopolymers are broken down to release the glucose monomers by either enzymatic or chemical means, in a process known as saccharification. This released glucose can be afterwards fermented for the production of bioethanol. However, the effectiveness of the saccharification step is restricted by several factors, such as the variability of the biomass, its inherent recalcitrance to digestion, as well as its heterogeneous molecular composition that includes, for instance, lignin polymers (5, 6). To overcome these hurdles, several physical and chemical pre-treatment methods are employed by biorefineries, such as: exposure to steam or liquid hot water; acidic treatments involving phosphoric, oxalic, acetic, or sulphuric acids (7, 8); and alkaline treatments, such as slaked lime or ammonia fiber explosion/expansion (AFEX) (6, 9–11). In our investigation, we focus on the interplay between the various factors which are pivotal in shaping the enzymatic saccharification dynamics of pre-treated biomass.

In addition to experimental studies, several modelling approaches have been developed at multiple scales, to understand the structure and recalcitrance properties of lignocellulose, and to decipher their role in determining saccharification yield. Each of these approaches brings its own set of advantages and disadvantages, which have been nicely summarised in the comprehensive review by Ciesielski et al. (12). At the lowest scale, the typically used methods include density functional theory (DFT), quantum mechanics/molecular mechanics (QM/MM), and molecular dynamics (MD) (13–16), which are used to address, for instance: pyrolysis (17, 18), the detailed structure and properties of lignocellulosic biomass (19), enzyme mechanisms (20, 21), and the effects of lignin binding on cellulose and cellulase enzymes (16). However, these methods are very expensive in terms of computational resources, and have limitations, e.g. being unable to depict biopolymers at the scale of seconds. To counter these disadvantages, other approaches have been used e.g. coarse-grained molecular dynamics with beads or pseudo atoms as elementary units (22). Besides, Kumar and Murthy employed further alternatives (i.e. Monte Carlo simulations) to study the enzymatic digestion of a cellulose bundle under the action of endoglucanase (EG), cellobiohydrolase (CBH), and *β*-glucosidase (BGL), by considering glucose molecules as the minimal substrate building blocks (23, 24). This model simulated a substrate of cellulose (including its crystallinity) along with hemicellulose and lignin. However, the impact of crystallinity on saccharification dynamics was not investigated, and strong discrepancies were observed between simulation results and experimental data. Instead, using Ordinary Differential Equations, Griggs et al. developed a mechanistic and kinetic model that simulated the action of a cellulase enzyme cocktail on a purely cellulosic substrate. It highlighted the enzymatic synergism, showed a good agreement with experimentally observed cellulose chain length distributions from literature, and displayed a semi-quantitative agreement with experimental saccharification time-course data (25, 26). However, the model contained significant simplifications, as the simulated substrate was purely consisting of cellulose. In addition, agent-based modelling approaches were employed by both Vetharaniam et al. (27) and Asztalos et al. (28) to highlight important phenomena. The former investigated enzymatic synergism, while considering hemicellulosic sugars and crystalline cellulose. The latter studied the synergistic action of multiple cellulases, while accounting for inter-chain hydrogen bond breaking, hydrolysis of glycosidic bonds, and adsorption and desorption of the cellulases on the substrate. Nonetheless, both of these studies contain several pitfalls, for instance, the omission of lignin in the substrate composition, or the consideration of a two-dimensional and purely cellulosic substrate. To fill these gaps, and build a comprehensive model of lignocellulose and its enzymatic saccharification dynamics, we introduced a stochastic biophysical model (29). This model did not only represent the three-dimensional structure of the lignocellulose material, the specific action of the enzymes, the crystallinity of cellulose and hemicellulose, and the role of lignin, but it also demonstrated an exceptional ability in quantitatively reproducing experimental saccharification time-course data.

Considering, on the one hand, the challenges of interpreting the experimental saccharification time-course data and the variability of lignocellulosic biomass, and on the other hand, the rapidly growing demand for renewable energy; it becomes apparent that the identification of the factors driving lignocellulose saccharification is needed in order to guide technological development efforts. To achieve this, in this study, we develop a full pipeline for the detailed Sobol’s sensitivity analysis (SSA) of the nineteen input parameters of our complex stochastic biophysical model (29). To consider realistic biorefinery processing lines, as well as high potential plant material for the production of biofuels, we focus on the illustrative case of corn stover under different pre-treatment conditions, and select two experimental datasets from the literature. They respectively deal with three different pre-treatment severities (30) and three different pre-treatment temperatures (31). We supplement our computational findings with analytical calculations, leading to a large reduction in the dimensionality of the parameter space, together with highlighting the key factors that drive the enzymatic digestion process.

## MATERIALS AND METHODS

### A Brief Reminder of our Stochastic Biophysical Model

Our stochastic biophysical model (29) utilises a Gillespie algorithm (32, 33) to simulate the enzymatic saccharification of a single lignocellulose microfibril. The three-dimensional structure of the microfibril is resolved at the level of monomers for the constituent biopolymers, i.e. glucose for cellulose, xylose for hemicellulose, and monolignols for lignin. As shown in figure. 1, the microfibril’s cellulosic core is surrounded by layers of hemicellulose and lignin polymers (34), and the crystalline properties of both cellulose and hemicellulose are taken into consideration. These are quantified by two parameters: i) the crystallinity fraction, which expresses the number of crystalline bonds over the total number of bonds of a certain type, for either cellulose or hemicellulose, and ii) the digestibility ratio, which describes how much harder it is to digest a crystalline bond as compared to its amorphous counterpart, for either cellulose or hemicellulose. The digestibility ratio is defined as:

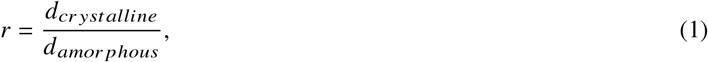

where *d*_*crystalline*_ and *d*_*amorphous*_ are the propensities for the digestion of a crystalline and an amorphous bond, respectively. We showed that the crystalline bonds are much harder for the enzymes to digest in comparison to the amorphous ones, such that *r*, that could lie between 0 (crystalline bonds cannot be digested) and 1 (crystalline and amorphous bonds cannot be distinguished), is typically in the range [10^−2^, 10^−3^].

**Figure 1:**
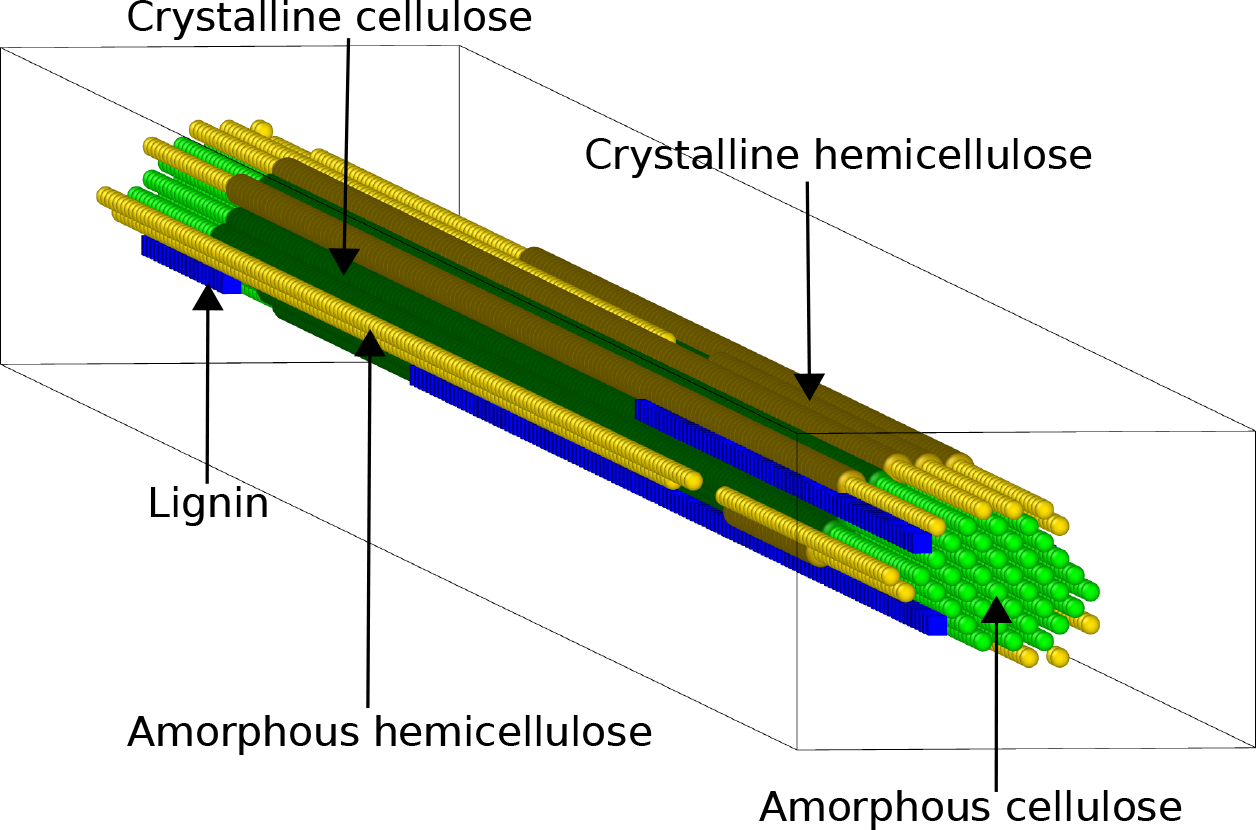
Schematic representation of the three-dimensional microfibril structure showing cellulose polymer chains at the core (in green), surrounded by outer layers of hemicellulose (in yellow) and lignin polymers (in blue). The crystalline regions for cellulose and hemicellulose are represented in darker shades of green and yellow, respectively.

The enzymatic digestion of the microfibril is carried out by an adjustable cocktail of cellulase and xylanase enzymes. Three classes of cellulase enzymes are defined in the model: endoglucanase (EG), cellobiohydrolase (CBH), and β-glucosidase (BGL). EG can digest any exposed bond on a cellulose polymer, with a reaction rate *K*_*EG*_, except the two outermost ones. CBH, a processive enzyme, attaches to the free ends of cellulose polymer chains at a rate 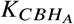; and processively cleaves off cellobiose units with a rate 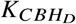. BGL completes the saccharification process by digesting the released cellobiose molecules into two glucose monomers, with a rate *K*_*BGL*_. Unlike cellulases, in our model, xylanase has a non-specific action such that it can digest any exposed hemicellulose bond, with a rate *K*_*XYL*_. The number of respective enzymes are noted *n*_*EG*_, *n*_*CBH*_, *n*_*BGL*_ and *n*_*XYL*_. To account for the impact of lignin on the saccharification process, we also consider the non-productive adsorption of enzymes on it, which we quantify by the parameter *Lignin adhesion rate*, that denotes the number of monolignols involved in binding a single enzyme.

This model (29) has been extended to also include the inhibition of cellulase enzymes by their end-products, i.e. cellobiose and glucose. Specifically, free cellobiose and glucose molecules bind to the cellulase catalytic sites and deactivate them; thus, reducing the effective number of cellulase enzymes capable of saccharification. The number of available enzymes (*n*_*y*_) is given by the general expression:

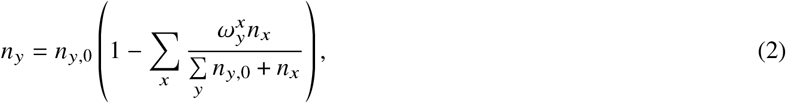

where *x* = *glc, cbs* (for glucose and cellobiose, respectively) and *y* = *EG, CBH, BGL*. The parameter 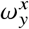 denotes the inhibition binding affinity of the inhibitor (*x*) on the enzyme (*y*), while *n*_*x*_ is the number of inhibitor molecules and *n*_*y*,0_ the number of enzymes if there was no inhibition present (i.e. like at the start of the simulation).

### Sobol’s sensitivity analysis

Using SSA allows us to highlight the model parameters having the highest impact on the dynamics of the enzymatic saccharification process (35, 36). We consider a function *f* with input parameters **x** ∈ ℝ^*p*^, where *p* is the dimension of the input parameter space, such that, in our case, *f* represents the absolute value of the difference between the simulated and the experimental saccharification curves and *p* = 19. For a specific point in the parameter space **x**^*^, the absolute value of the difference between the simulated and the experimental saccharification curves is denoted as Y = *f* (**x**^*^). SSA is a variance-based measure of global sensitivity such that the input parameter x_i_ with the highest impact is the one that most contributes to the variance of *f*. The total Sobol’s index (*S*_*T*_) of the input parameter x_i_ measures the contribution of x_i_ to the variance of *f*, including all contributions due to its interactions, of any order, with any other input parameter.

To perform our analysis, we use the open-source Python library SALib (37, 38) that typically follows four steps. We construct a pipeline to connect our previously published stochastic biophysical model of lignocellulose (29) to the SALib library, such that it: i) selects the parameters whose influence will be analysed; ii) runs the *sample* function of SALib to generate the list of input parameter sets, depending on the number of parameters to be analysed; iii) runs the model for each parameter set and calculates the absolute value of the difference between the simulated and the experimental saccharification curves; iv) runs the *analyze* function of SALib on the calculated absolute differences to determine the variances, and hence, the Sobol’s indices (37, 38). Our large amount of input parameters (i.e. *p* = 19) requires a high number of parameter sets to be tested for each pre-treatment condition (i.e. 21, 504). Since the model performs stochastic simulations we additionally need to repeat each run 10 times to obtain a single simulated saccharification curve. Thus, the analysis of the three pre-treatment conditions for each of the two experimental datasets that we consider here sums up to a total of 1, 290, 240 simulation runs. We therefore limit ourselves to studying the total Sobol’s indices (*S*_*T*_).

## RESULTS

### Impactful parameters according to Sobol’s sensitivity analysis

Figure. 2 shows the results of the SSA (i.e. the total indices, noted *S*_*T*_) performed on the nineteen parameters of our stochastic biophysical model comparing to the experimental datasets of Bura et al. (30) and Liu et al. (31), that each contains three pre-treatment conditions. These indices highlight the model parameters with the highest impact on the saccharification time-courses. It is important to note that Sobol’s indices may only be evaluated relatively within the results of a single sensitivity analysis. Hence, even for analyses with similar functions and input parameters, the results cannot be compared as absolute values. Here, for each pre-treatment condition, we choose to scale the *S*_*T*_ values with respect to the corresponding total Sobol’s index of the CBH reaction rate, i.e. 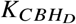. Consistently across all the six conditions considered, the same five parameters play the most significant roles in determining the saccharification dynamics. These can be classified in two groups: i) the three parameters relating to the activity of CBH, i.e. the CBH processive reaction rate 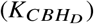, the inhibition binding affinity of glucose to CBH 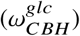, and the initial number of CBH enzymes in the system (*n*_*CBH*_); ii) the two parameters quantifying the crystallinity properties of cellulose, i.e. the digestibility ratio (*r*) and the cellulose crystallinity fraction (noted *X*). Interestingly, although the crystallinity of cellulose plays a major role in determining the saccharification dynamics, that is not the case for hemicellulose’s crystallinity. This effect can be explained by the fact that hemicellulose is almost entirely removed from the substrate during pre-treatment (30, 31), and because the time-courses considered here are those of glucose conversion.

**Figure 2:**
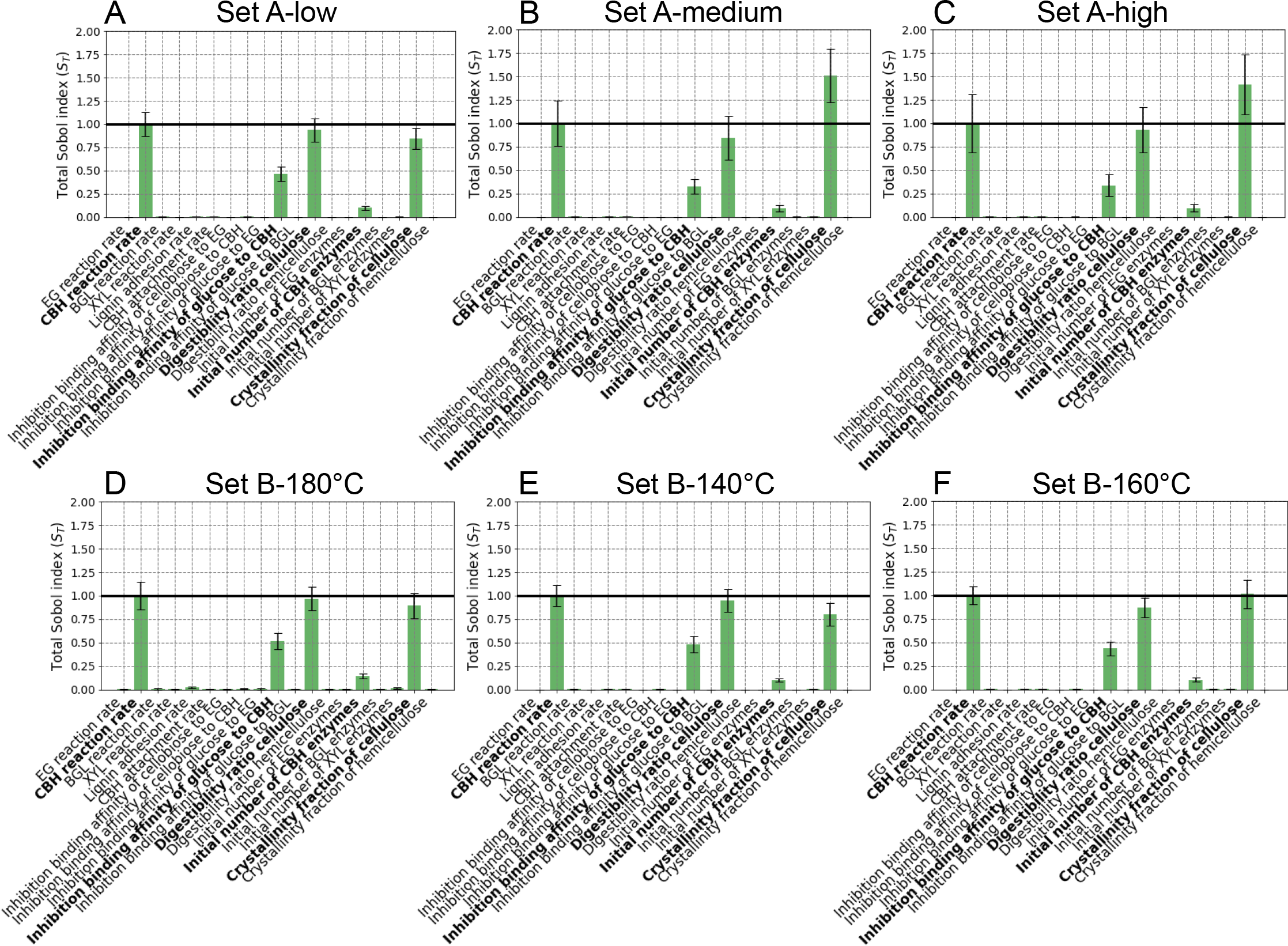
Total Sobol’s indices (*S*_*T*_) obtained when our stochastic biophysical model is compared to the experimental saccharification time-courses of Bura et al.’s data (30) (three different pre-treatment severities: A low; B medium; and C high) and of Liu et al.’s data (31) (three different pre-treatment temperatures: D 140°C; E 160°C; and F 180°C). The indices are normalised with respect to the total Sobol’s index (*S*_*T*_) of the CBH processive reaction rate 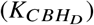 of each pre-treatment condition, which is indicated with a black line. The columns from left to right show the conditions resulting in increasing final saccharification yield (at 72 hours), for each dataset (see table 1).

Amongst all total Sobol’s indices, the one of the cellulose crystallinity fraction shows the strongest variation across pre-treatment conditions. Particularly for Bura et al.’s data (Set A), its value increases going from low to medium severities, and then drops for high severity. For Liu et al.’s data (Set B), although some minor variation in the *S*_*T*_ of the cellulose crystallinity fraction can be observed amongst the pre-treatment conditions, those are not significant. In addition, we observe that this Sobol’s index is strongly correlated to the glucose yield after 72 hours when considering all six conditions, as shown in figure. 5A.

**Table 1:** Final saccharification yield (at 72 hours), crystallinity fraction of cellulose for the best fit, and *S*_*T*_ of cellulose crystallinity fraction, for Bura et al.’s data (30) (Set A, three different pre-treatment severities: low; medium; and high) and for Liu et al.’s data (31) (Set B, three different pre-treatment temperatures: 180°C; 140°C; and 160°C).

| Set | A |  |  | B |  |  |
| --- | --- | --- | --- | --- | --- | --- |
| Pre-treatment condition | low | medium | high | 180°C | 140°C | 160°C |
| Glucose conversion at 72 h | 58.2 | 87.5 | 100 | 33.02 | 52.36 | 65.25 |
| Crystallinity fraction of cellulose for the best fit | 0.50 | 0.23 | 0.09 | 0.72 | 0.59 | 0.41 |
| $S_T$ of cellulose crystallinity fraction | 0.847 | 1.512 | 1.416 | 0.894 | 0.799 | 1.015 |

### Effect of cellulose crystallinity

Figure. 3 shows a scatter plot of the absolute value of the difference between the simulated and the experimental saccharification time-courses (noted Y) *versus* the cellulose crystallinity fraction for the six pre-treatment conditions considered. Each point represents a specific set of input parameters. When Y is less than 150, the points are colour-coded in blue. When it is bigger than 150, if the simulated curve is above the experimental one, the points are green, otherwise, they are turquoise. For a given pre-treatment condition, the number of turquoise points increases with the cellulose crystallinity fraction (moving from left to right along the x-axis), showing that the simulated saccharification curve lies below the experimental one; and thus, that the glucose conversion through time diminishes as the cellulose crystallinity fraction increases. Consistently, when instead the cellulose crystallinity fraction decreases (moving from right to left along the x-axis), almost exclusively green points are found, showing that the simulated saccharification curve lies above the experimental one; and thus, that the glucose conversion through time increases as the cellulose crystallinity fraction diminishes. It should be noted that the maximum value of Y at which the green points saturate, reflects the parameter sets responsible for the fastest possible digestion scenario, for a specific value of the cellulose crystallinity fraction. Analogously, the turquoise points are limited by the case of the slowest digestion throughout the entire time-course, for a specific value of the cellulose crystallinity fraction (not visible on B and C because of the displayed y-axis range). For each of both the datasets, across pre-treatment conditions, with increasing final saccharification yields (columns from left to right), the maximum value of Y for the green points decreases while that of the turquoise points increases. This can be simply explained by the areas of the corresponding zones, respectively above and below the experimental saccharification time-courses. The faster the saccharification, the higher the experimental curve, the more area is turquoise and the less is green, and *vice versa*. This is confirmed when comparing the maximum values of Y for the green points across the two datasets. Ranking the pre-treatment conditions on the basis of these maximum values of Y in the decreasing order, corresponds to the increasing order of the final conversion yields at 72 hours (see table 1):

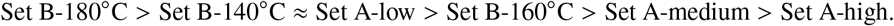

**Figure 3:**
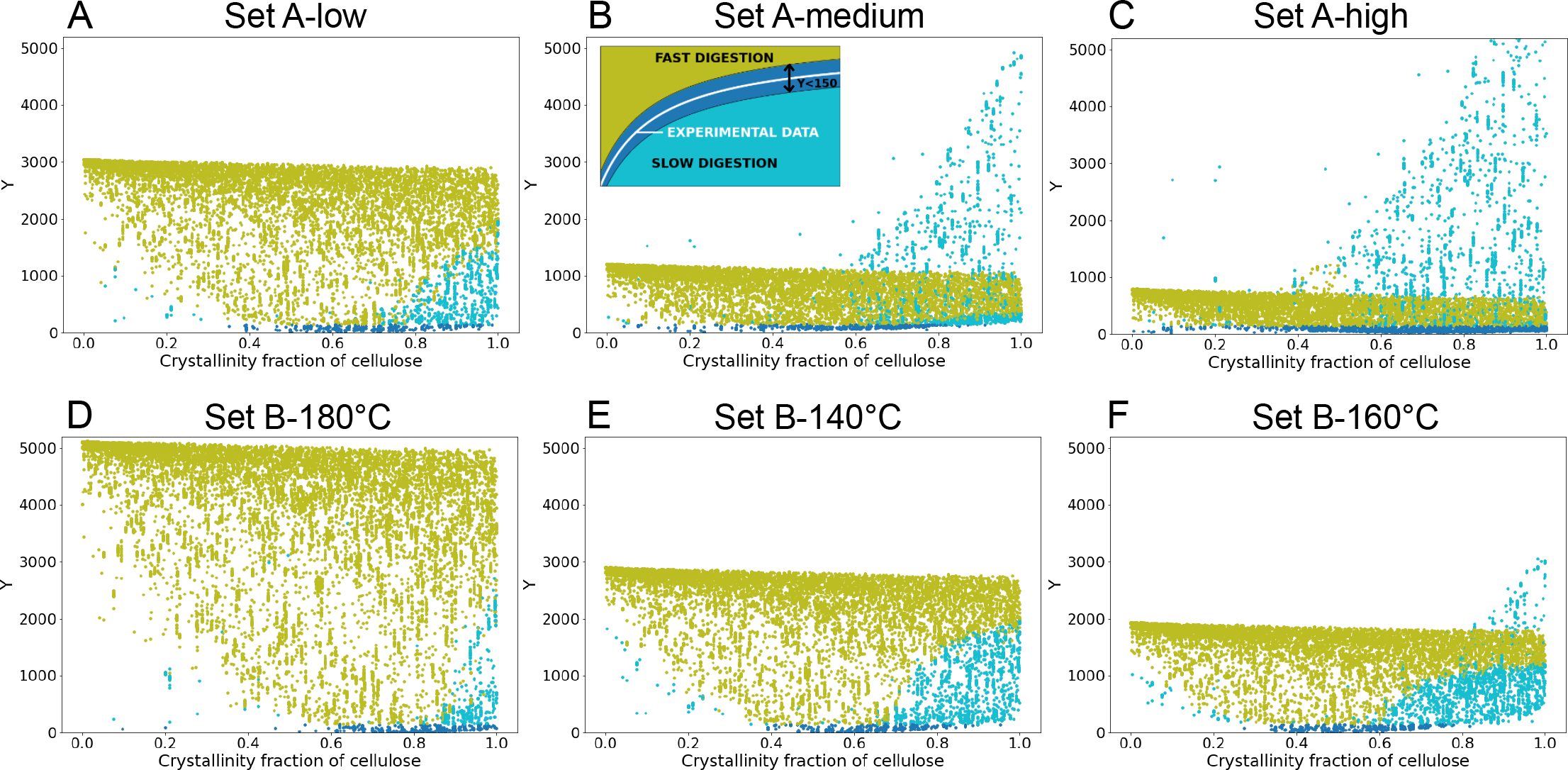
Absolute value of the difference between the simulated and the experimental saccharification time-courses (Y) *versus* the cellulose crystallinity fraction for Bura et al.’s data (30) (three different pre-treatment severities: A low; B medium; and C high) and for Liu et al.’s data (31) (three different pre-treatment temperatures: D 140°C; E 160°C; and F 180°C). The columns from left to right show the conditions resulting in increasing final saccharification yield (at 72 hours), for each dataset (see table 1). Points are colour-coded according to the inset in B.

In figure. 4, for each pre-treatment condition, we specifically focus on the parameter sets that closely reproduce the experimental saccharification time-courses (blue points with Y < 150), and highlight the best fit with a red circle. The spread of the blue data points is evaluated at Y = 75, by considering 98% of the points. For Liu et al.’s data (Set B), this spread shows very little variations around ca. 0.32, while, for Bura et al.’s data (Set A), this quantity significantly increases with pre-treatment severity (columns from left to right). Overall, for both datasets, we observe that the spread of the blue points closely matching the experimental saccharification time-courses is much narrower for Liu et al.’s data (Set B) than for Bura et al.’s data (Set A).

**Figure 4:**
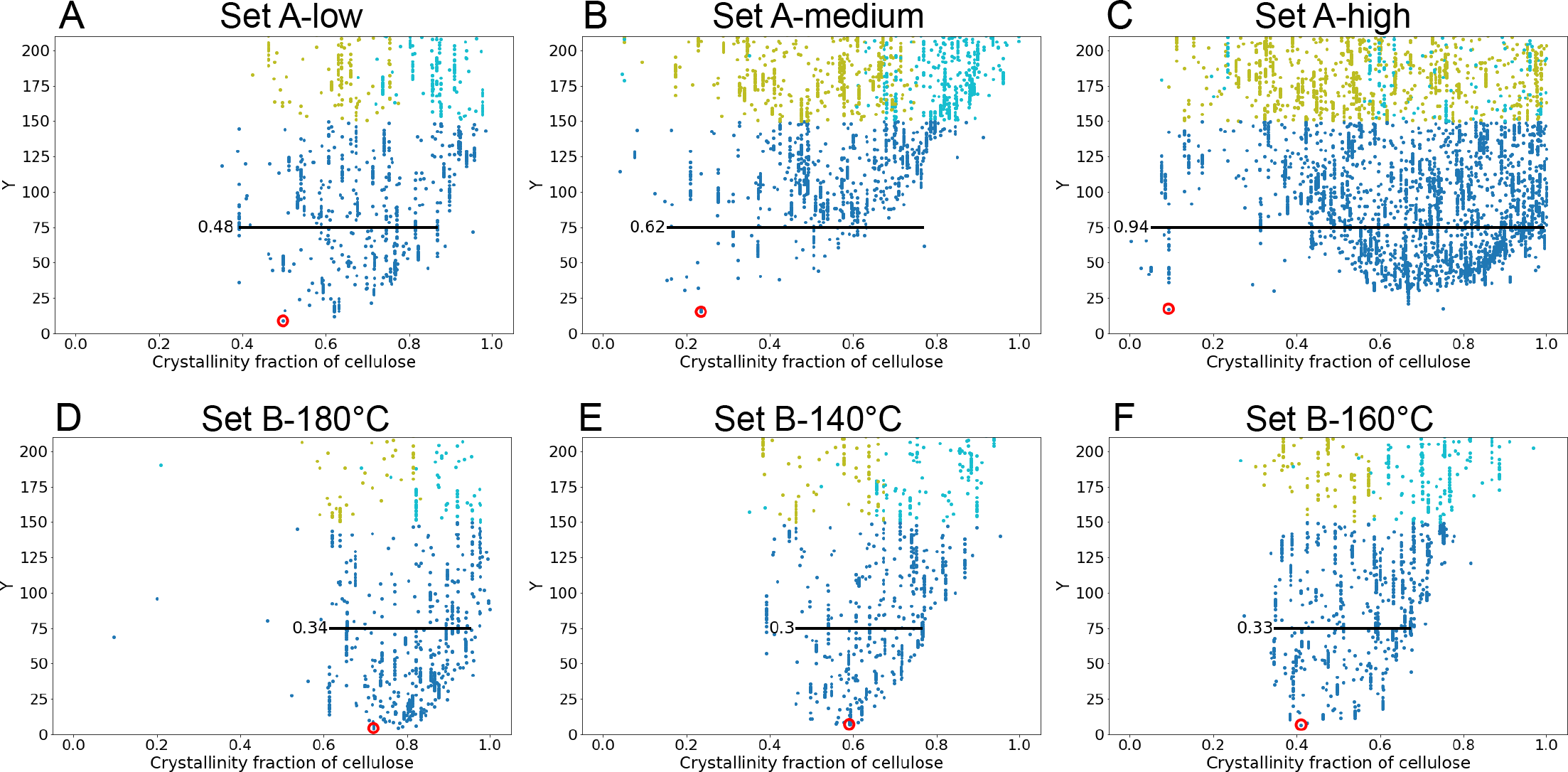
Zoomed in view of the absolute value of the difference between the simulated and the experimental saccharification time-courses (Y) *versus* the cellulose crystallinity fraction for Bura et al.’s data (30) (three different pre-treatment severities: A low; B medium; and C high) and for Liu et al.’s data (31) (three different pre-treatment temperatures: D 140°C; E 160°C; and F 180°C). The columns from left to right show the conditions resulting in increasing final saccharification yield (at 72 hours), for each dataset (see table 1). Points are colour-coded according to the inset in figure. 3B. For each pre-treatment condition, the red circled point corresponds to the lowest value of Y, i.e. the best fitting parameter set. The black line indicates the spread of the distribution below the threshold Y = 75, including 98% of the points (excluding 1% to the left and 1% to the right).

**Figure 5:**
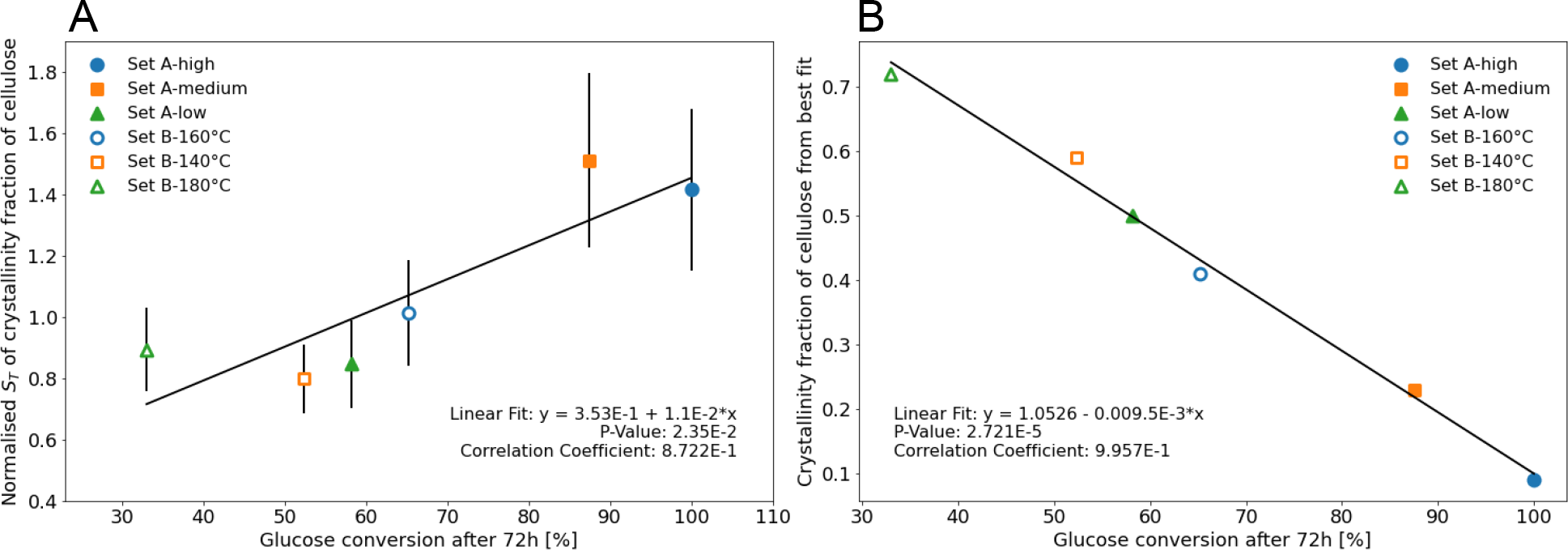
A: Normalised *S*_*T*_ of the cellulose crystallinity fraction *versus* final glucose conversion yield (at 72 hours) for the six pre-treatment conditions considered shows a positive correlation. B: Crystallinity fraction of cellulose for the best fits *versus* final glucose conversion yield (at 72 hours) for the six pre-treatment conditions considered shows a negative correlation.

It means that as the cellulose crystallinity fraction (x-axis) varies, the value of Y is more impacted in the Set B as compared to in the Set A. In Liu et al.’s data (Set B), the blue points with the lowest Y values also collectively converge towards the best fit (red circle), which is not the case for Bura et al.’s data (Set A). Noticeably, in the case of high pre-treatment severity for Bura et al.’s data (C), a large number of points are clustered for values of cellulose crystallinity fraction between 0.45 and 1, while the outlying best fit shows a value of ca. 0.10. This observation indicates that when designing a fitting algorithm, one must carefully navigate the parameter space since seemingly good fits can return very inaccurate parameter values, even for the most impactful parameter of the system. Additionally, it stresses the importance of a critical assessment of the feasibility of the obtained best fit parameter values. In figure. 4, across pre-treatments, the cellulose crystallinity fraction of the best fits (red circled) decreases as the glucose conversion after 72 hours increases (columns from left to right), consistently with the reported role of crystallinity in boosting saccharification recalcitrance. Overall, as illustrated in figure. 5, on the one hand, the cellulose crystallinity fraction of the best fits shows a strong negative correlation with the glucose conversion at 72 hours; while, on the other hand, the normalised *S*_*T*_ of the cellulose crystallinity fraction has a positive correlation with the glucose conversion at 72 hours.

Figure. 6 displays the time-courses of the best fits (dashed lines) and in case the value of the cellulose crystallinity fraction is either increased or decreased by 0.05 (solid lines), for both Bura et al.’s and Liu et al.’s datasets. From the area bounded by the respective solid lines through time, it clearly arises that a variation in the cellulose crystallinity fraction impacts the later times (plateau region) much stronger than it does for the earlier times (region with a steeper slope). This observation is consistent with the amorphous bonds being digested first. Moreover, the area of the bounded region between the solid lines is greater for Liu et al.’s pre-treatment conditions as compared to the ones of Bura et al.’s. This can easily be attributed to the profiles of the respective saccharification curves, since the duration of the plateau region is in general longer for Liu et al.’s experimental data. Our observation that in close vicinity to the best fit, variations in the cellulose crystallinity fraction for Liu et al.’s data, lead to stronger deviations from the best fits, as compared to the case of Bura et al.’s data, is also consistent with the spread of the blue points at Y = 75 in figure. 4 being smaller for Liu et al.’s data.

**Figure 6:**
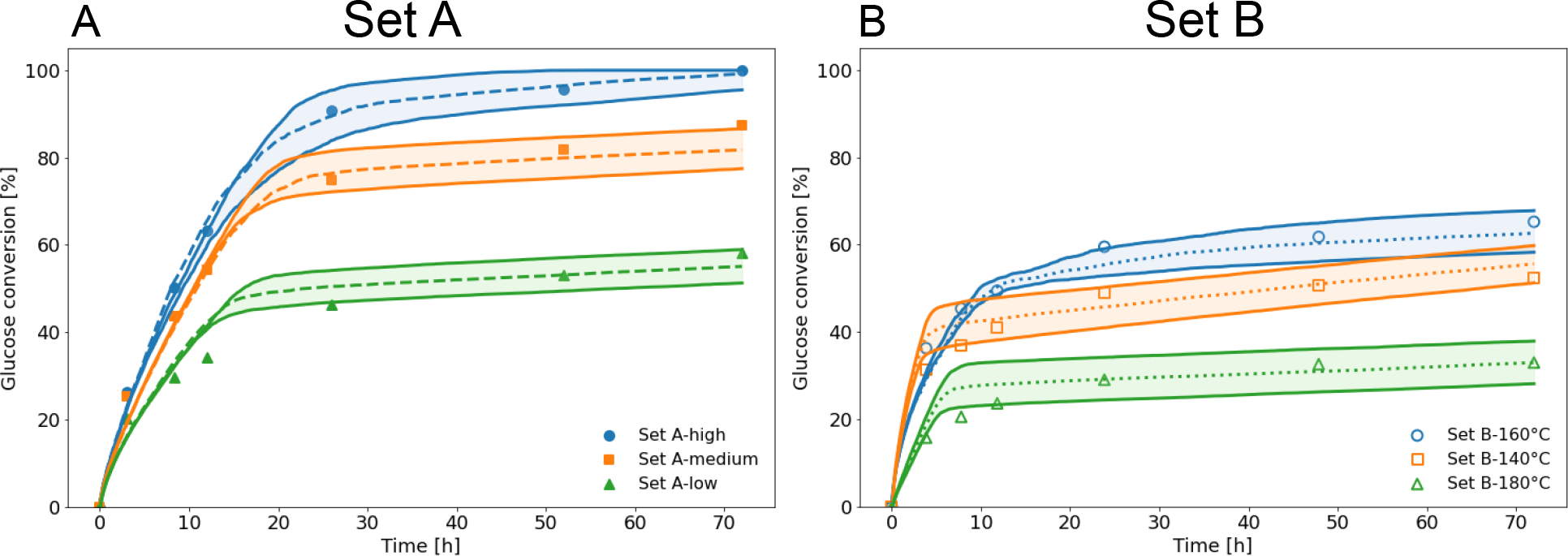
Experimental saccharification time-courses (points) for A: Bura et al.’s data (Set A) (30), and B: Liu et al.’s data (Set B) (31). The best fits (corresponding to red circles in figure. 4) are shown as dashed lines, while solid lines depict simulated saccharification curves if the value of the cellulose crystallinity fraction is either increased or decreased by 0.05 from the best fitted value. The area bounded by the respective solid lines is equal to: Set A-high: 510.21, Set A-medium: 517.50, Set A-low: 432.13, Set B-160 °C: 485.46, Set B-140 °C: 628.94, and Set B-180 °C: 664.52 (the area unit is hours).

### Role of CBH

To decipher the specific influence of the CBH enzyme, we consider a fully amorphous cellulose microfibril, without any hemicellulose or lignin. This simplification of the substrate allows us to focus on the role of CBH by omitting the impact of crystallinity, in addition to that of lignin and hemicellulose. We call this substrate configuration the *test sample*.

In Figure. 7A, each point depicts the absolute value of the difference between the saccharification time-courses simulated for a specific set of input parameters and for the *test sample versus* the CBH reaction rate. We clearly observe the clustering of the green points that indicate faster digestion than the *test sample*. This cluster is bounded by a radical function-like shape, which, for a fixed CBH reaction rate, denotes the input parameter set responsible for the fastest possible digestion scenario. As the CBH reaction rate increases (moving from left to right along the x-axis), the digestion gets faster; thus it is only logical to note that the maximum value of Y for the green points increases. This is consistent with the clustering at the upper boundary of this radical function-like shape of the dark purple points in figure. 7B and of the yellow ones in figure. 7C. For a particular CBH reaction rate, these correspond to low inhibition binding affinity of glucose to CBH and to a high initial number of CBH enzymes, respectively, both of which contribute to increasing the CBH propensity. From figure. 7A, we observe that the number of turquoise points indicating slower digestion than that of the *test sample*, decreases with increasing CBH reaction rate. Similar to our remark for figure. 3, the upper limit of the turquoise points is not visible due to the choice of the y-axis range. We observe that for a fixed value of Y, the turquoise points of figure. 7A, show a colour shift from dark purple to light yellow moving from left to right along the x-axis in figure. 7B, whereas the reverse tendency is noted in figure. 7C. In other words, increasing the inhibition binding affinity of glucose to CBH or decreasing the initial number of CBH enzymes both contribute to the decrease of the CBH propensity, which is nonetheless balanced by an increase in CBH reaction rate, thus resulting in an unchanged value of Y. Finally, in figure. 7A, the blue points (Y < 150), that expectedly lie at the bottom of the plot, display the same colour shifts along the x-axis of figure. 7B and figure. 7C as described above for the turquoise points. Interestingly, although their distribution is denser at low values of CBH reaction rate, the lowest Y value, which is obtained for the input parameter set best fitting the *test sample*, is observed at ca. 45 reactions per hour, which corresponds to both a high inhibition binding affinity of glucose to CBH and to a low initial number of CBH enzymes. Both for the blue and the turquoise points of figure. 7A, the colour shifts observed are more prominent in figure. 7B than in figure. 7C, which can be attributed to a higher total Sobol index (*S*_*T*_) for the inhibition binding affinity of glucose to CBH (figure. 7B) as compared to that of the initial number of CBH enzymes (figure. 7C). For the other two cellulases (EG and BGL), the absolute value of the difference between the saccharification time-courses simulated for a specific set of input parameters and for the *test sample versus* the respective enzyme reaction rate reveals no impact of the enzyme reaction rate (see Supplementary Material).

**Figure 7:**
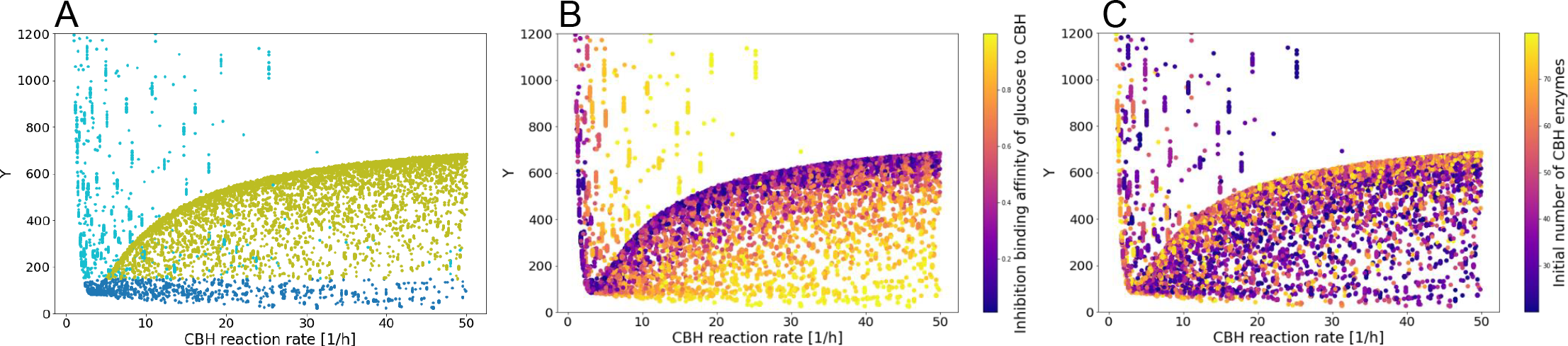
Absolute value of the difference between the saccharification time-courses simulated for a specific set of input parameters and for the *test sample versus* the CBH reaction rate (i.e. the CBH parameter with the highest *S*_*T*_ index). The colour-code indicates: A the relative position of the simulated saccharification curve with respect to the one of the *test sample*, as defined in the inset of figure. 3B; B the inhibition binding affinity of glucose to CBH (i.e. the CBH parameter with the second highest *S*_*T*_ index); and C the initial number of CBH enzymes (i.e. the CBH parameter with the third highest *S*_*T*_ index). The CBH reaction rate is sampled within the range [1, 10^4^] reactions per hour. However, it is only displayed in the range [0, 50] where significant variations can be observed.

**Figure 8:**
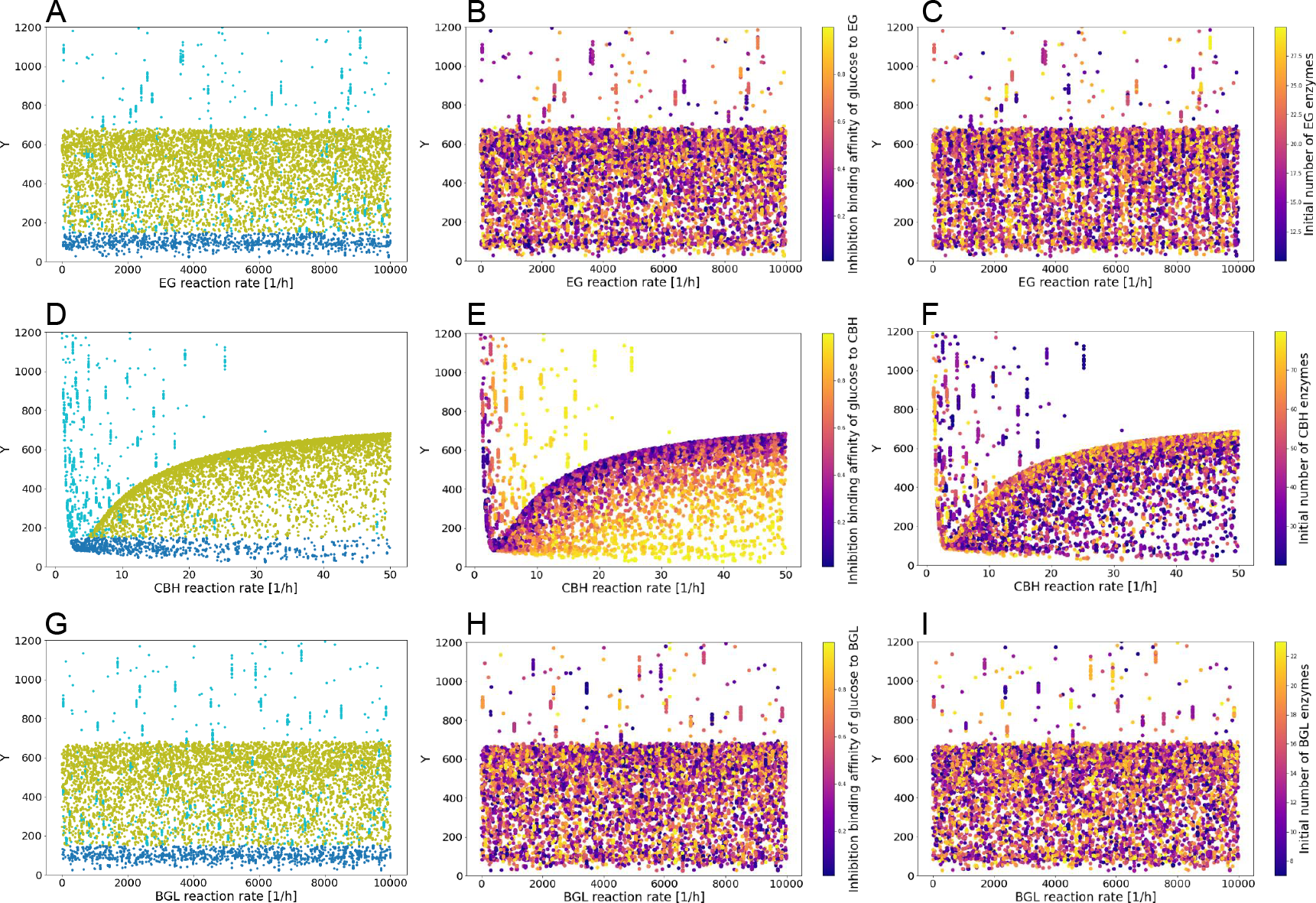
Absolute value of the difference between the saccharification time-courses simulated for a specific set of input parameters and for the *test sample versus* the enzyme reaction rate of EG: A, B, and C; CBH: D, E, and F; and BGL: G, H, and I. The colour-code indicates: in A, D, and G: the relative position of the simulated saccharification curve with respect to the one of the *test sample*, as defined in the inset of figure. 3B; in B, E, and F: the inhibition binding affinity of glucose to the respective cellulase; and in C, F, and I: the initial number of cellulase enzymes of the respective type.

## ANALYTICAL CALCULATIONS

In this section, we use a simplified system and some elementary calculations to confirm our computational results and to determine their range of validity for different parameter values. For this purpose, we consider a system that consists of a pure cellulose polymer being digested by a cocktail of cellulases, comprising a number of EG, CBH, and BGL enzymes. The process starts with the polymer having a degree of polymerisation (DP) equals 2*N* (noted *DP*_2*N*_), and continues until 2*N* glucose molecules have been released. The propensity for the diffusive action of an enzyme (*y*) is given as:

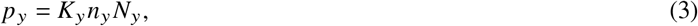

where *K*_*y*_ is the reaction rate of the enzyme *y* (i.e. digestion in case of a diffusive enzyme, and attachment in case of a processive enzyme), *n*_*y*_ is the number of molecules of enzyme *y*, and *N*_*y*_ is the number of possible attack points (or reaction sites) for the specific enzyme on that substrate. The characteristic time for such a reaction to occur can be simply expressed as:

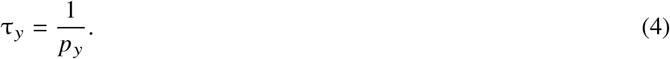

The digestion of the polymer of length *DP*_2*N*_ proceeds according to the following steps.

### EG reaction

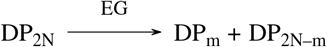

The polymer is cut by EG into 2 pieces of respective lengths *DP*_*m*_ and *DP*_2*N* −*m*_. For ease of illustration, we can consider *m* = *N* without any loss of information.

### CBH reaction

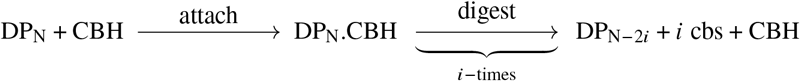

The reaction by CBH has two steps, first the attachment of CBH to one of the two free ends of any polymer, followed by the processive digestion steps, each releasing cellobiose.

### BGL reaction

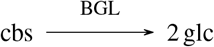

Cellobiose is digested by BGL to release 2 glucose molecules.

Considering an initial polymer in which all the bonds are amorphous, and using equations (3) and (4), we express the characteristic times for the digestion reactions by the cellulases in table 2. Let us now consider that the polymer has a cellulose crystallinity fraction of *X* (0 ≤ *X* ≤ 1) and a cellulose digestibility ratio of *r* (0 < *r* < 1). The characteristic time for all the bonds of this polymer to be degraded by an enzyme *y*, is the weighted sum of the characteristic times for the digestion of both the crystalline and the amorphous parts, and can be expressed as:

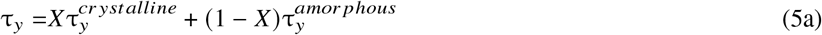

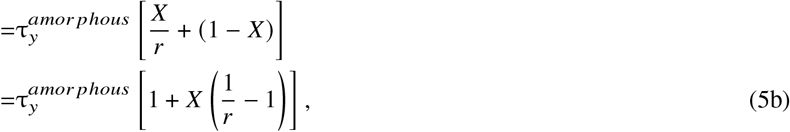

with 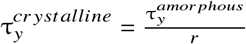. Using equation (5b), we can write the characteristic time for the reaction by each of the three cellulases as:

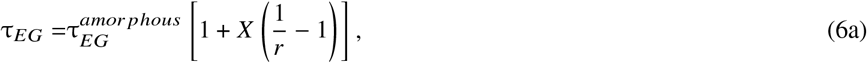

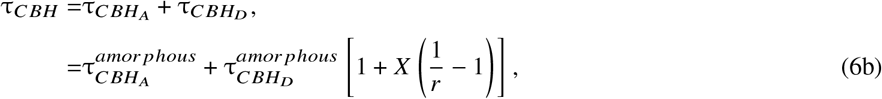

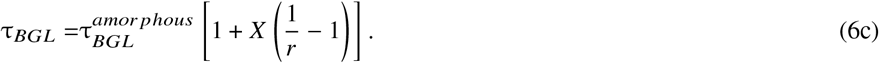

**Table 2:** Characteristic times for cellulase reactions on an amorphous cellulose chain of length *DP*_2*N*_.

| Reaction | Characteristic time | Comment |
| --- | --- | --- |
| EG | $\tau_{EG}^{amorphous} = \frac{1}{K_{EG}n_{EG}(2N-4)}$ | (2N-4) possible attack points for EG on a cellulose chain of $DP_{2N}$ |
| CBH attachment | $\tau_{CBH_A}^{amorphous} = \frac{1}{2 \times 2 \times K_{CBH_A} \times n_{CBH}}$<br>$= \frac{1}{4K_{CBH_A}n_{CBH}}$ | Two polymers of $DP_N$ have a total of 4 free ends |
| CBH digestion | $\tau_{CBH_D}^{amorphous} = 2 \times \frac{N}{2} \times \frac{1}{K_{CBH_D}}$<br>$= \frac{N}{K_{CBH_D}}$ | CBH reacts N/2 times on each polymer of $DP_N$ |
| BGL | $\tau_{BGL}^{amorphous} = N \times \frac{1}{n_{BGL}K_{BGL}N}$<br>$= \frac{1}{n_{BGL}K_{BGL}}$ | BGL reacts N times on N cellobiose molecules from a polymer of $DP_{2N}$ |

It should be noted that *r* is the *digestibility* ratio that discriminates between amorphous and crystalline bonds, and hence, it does not affect CBH’s attachment step, such that:

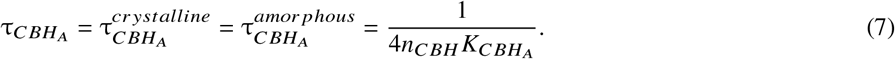

Substituting the values from table 2 into equation (6), we obtain:

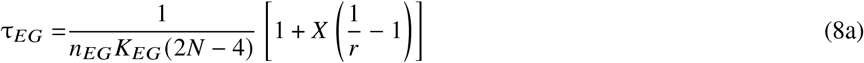

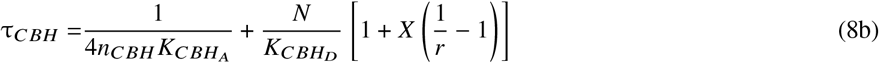

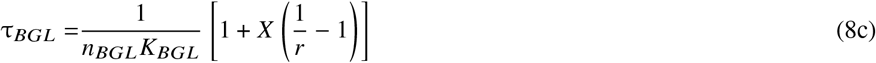

The end product inhibition of cellulases by glucose and cellobiose molecules is also known to affect the saccharification yield, by reducing the effective number of enzymes available for digestion. In our model, the effective number of enzymes (*n*_*y*_) can be expressed by the general expression:

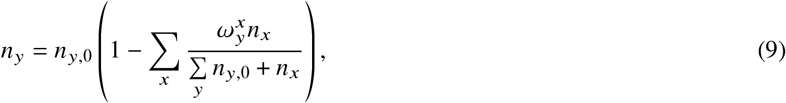

where *x* = *glc, cbs* and *y* = *EG, CBH, BGL*. The parameter 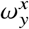 denotes the inhibition binding affinity of the inhibitor (*x*) on the enzyme (*y*), while *n*_*x*_ is the number of inhibitor molecules, and *n*_*y*,0_ the number of enzymes if there was no inhibition present (i.e. like at the start of the simulation). In our simulated system, we do not observe much effect of inhibition by free cellobiose, as it does not accumulate, due to the presence of a sufficient number of BGL enzymes. Thus, we can simplify the expression inequation (9) as:

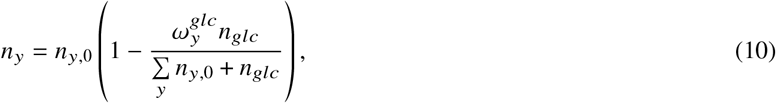

Substituting, equation (10) into equation (8) we obtain:

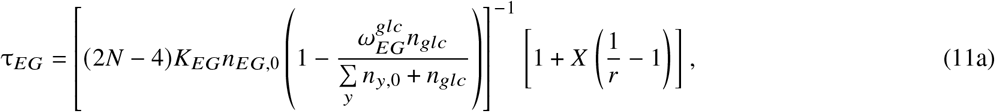

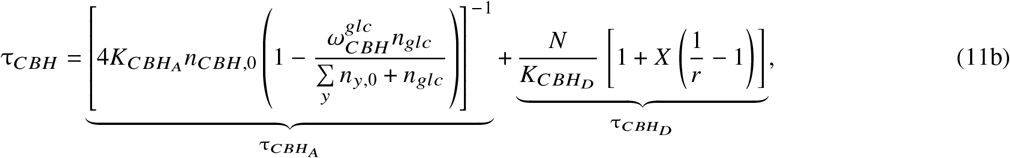

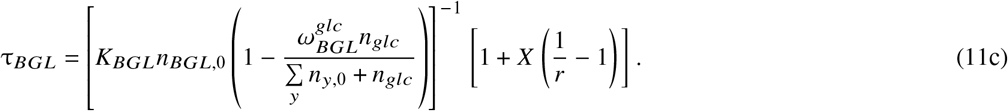

Since the characteristic time is the average duration a reaction needs to take place, the higher the characteristic time the slower that reaction. Thus, the limiting time step is that with the largest characteristic time. From our computational results, we know that the CBH reaction plays the key role in determining the saccharification yield. To support this, we need to prove that the characteristic time for the CBH reaction is higher than those for both EG’s and BGL’s reactions, i.e. that:

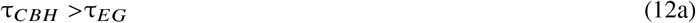

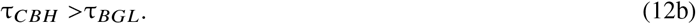

Since 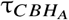 is always positive, it is sufficient to show that:

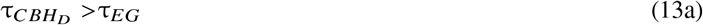

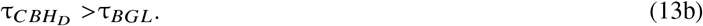

Thus, substituting the values from equation (11) into the inequations 13, we obtain:

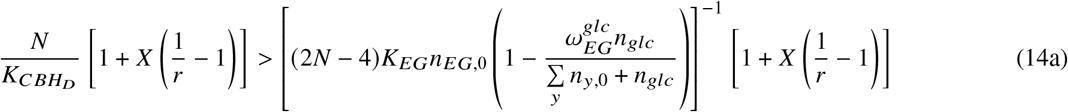

and

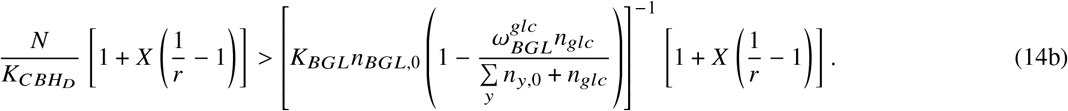

From the inequations 14a and 14b, we can cancel out the common factor 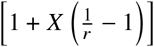 from both sides (having 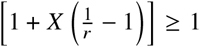 since (0 ≤ *X* ≤ 1) and (0 < *r* < 1)). We obtain:

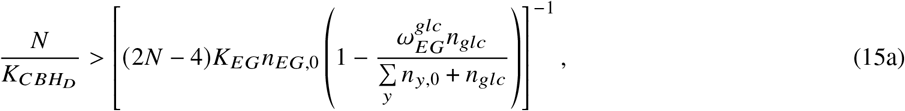

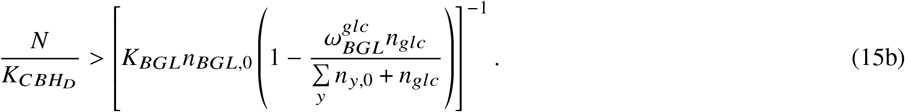

Both inequations are most constrained in the case when *n*_*EG*,0_ = *n*_*BGL*,0_ = 1, 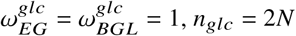, and 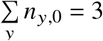.

Using the most constrained case, for the EG reaction, from inequation 15a, we can write for a polymer of *DP*_2*N*_ :

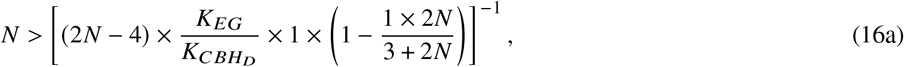

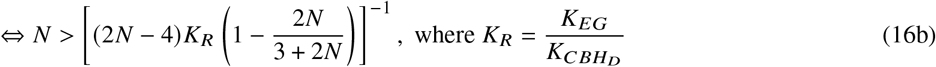

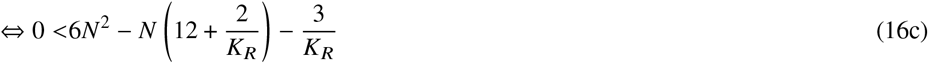

The second order polynomial *P*(*N*) on the R.H.S of inequation 16c, admits 2 real roots (*N*_1_ and *N*_2_), that can be written as:

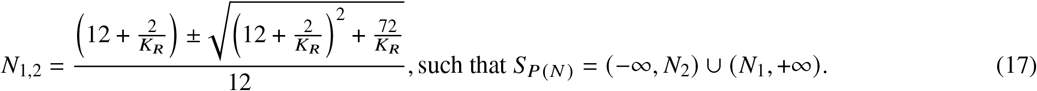

We study the limits of *N*_1_ and *N*_2_, and thus the validity of inequation 16c, depending on the value of *K*_*R*_ ∈ (0, +∞):

i. If

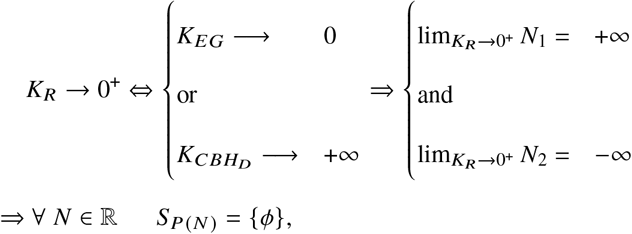

then inequation 16c is invalid for all *N* ∈ [3, +∞)
ii. If 0 < *K*_*R*_ < 1, then inequation 16c is valid only for the values of *N* that respect *S*_*P*(*N*)_ = (−∞, *N*_2_) ∪ (*N*_1_, +∞) depending on the values of *N*_1_ and *N*_2_ defined inequation (17).
iii. If *K*_*R*_ = 1, *N*_1_ = 2.53 and *N*_2_ = −0.19, then inequation 16c is valid for all *N* ∈ [3, +∞).
iv. If *K*_*R*_ → +∞, *N*_1_ → 2 and *N*_2_ → 0^+^, then inequation 16c is valid for all *N* ∈ [3, +∞).

Using the most constrained case, for the BGL reaction, from inequation 15b, we can write for a polymer of *DP*_2*N*_ :

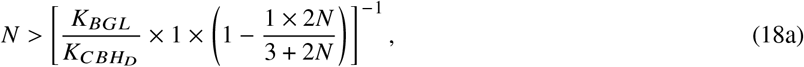

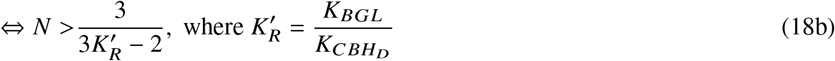

i. If 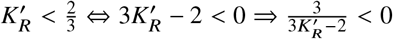, then inequation 18 is valid for all *N* ∈ [3, +∞).
ii. If 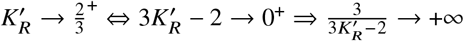, then inequation 18 is invalid for all *N* ∈ [3, +∞).
iii. If 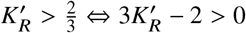, then inequation 18 is valid only for the values of *N* that respect 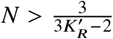.

The validity of the ineqations 16 and 18 for the aforementioned conditions on *K*_*R*_, 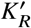 and *N* proves that the CBH reaction is the time-limiting step for such conditions.

For our model to best fit the experimental saccharification time-courses for the six pre-treatment conditions considered, the typical ranges of the enzyme kinetic rates must be 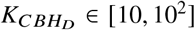, *K*_*EG*_ ∈ [10^2^, 10^3^], and *K*_*BGL*_ ∈ [10^2^, 10^3^]. Since for these values of enzyme kinetic rates, *K*_*R*_ ∈ [1, 10^2^] and 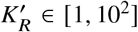 (and we also have *N* >> 3), our analytical approach confirms that CBH is the time-limiting step. As a consequence, it is only logical that the Sobol’s sensitivity analysis showed that CBH is the enzyme with the strongest impact.

Naturally, the parameters having the main impact on the saccharification yield are those that appear in the expression of τ_*CBH*_, i.e. inequation (11b). Since it has been sufficient to demonstrate that 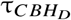 is greater than τ_*EG*_ and τ_*BGL*_ to show that the CBH reaction is the time-limiting step, we deduce that the parameters appearing in the expression of 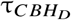 are the most important ones (see equation (11b)). These are: the CBH reaction rate 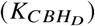, the cellulose crystallinity fraction (*X*), and the digestibility ratio of cellulose (*r*). The parameters appearing in the expression of 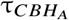 (see equation (11b)) have a somewhat lower impact. These are: the inhibition binding affinity of glucose to CBH 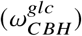, the CBH attachment rate 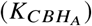, and the initial number of CBH enzymes (*n*_*CBH*,0_). The parameters 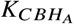 and *n*_*CBH*,0_ have a symmetric role in the expression of 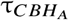. However, their typical value ranges are different by one order of magnitude (i.e. *n*_*CBH*,0_ ∈ [10, 10^2^], and 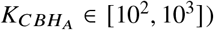). As a consequence, a deviation by one unit in *n*_*CBH*,0_ changes 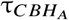 by a maximum of ca. 10%, whereas such a deviation for 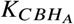, impacts 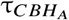 by at most 1%. This explains why the normalised total Sobol’s sensitivity index (*S*_*T*_) observed in figure. 2 for *n*_*CBH*,0_ is much higher than that of 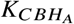.

Overall, our calculations show that there are only five impactful parameters. The ones with the highest importance are: the CBH reaction rate 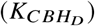, the cellulose crystallinity fraction (*X*), and the digestibility ratio of cellulose (*r*); while the second most influential ones are: the inhibition binding affinity of glucose to CBH 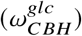, and the initial number of CBH enzymes (*n*_*CBH*,0_). These parameters and their grouping by impact strength are in full agreement with our computational results.

## CONCLUSION

The utilisation of crop residues from agricultural waste as an abundant and renewable raw material is a vivid research topic owing to its high potential for industrial applications, such as the production of bio-ethanol (3). The enzymatic extraction of simple sugars, for instance for their further downstream fermentation, faces several challenges, not only due to the variability of the biomass depending on its source, but also because of its complex structure and its interplay with several enzymes during the saccharification process, which eventually results in biomass recalcitrance (6, 39). To counter this effect and increase glucose conversion, several pre-treatment methods have been designed (5). Nonetheless, it was so far unclear which biophysical parameters of the system drive the saccharification process, despite their importance for the optimal design of experimental procedures, as well as to support the rationalisation of complex experimental datasets.

To fill this gap, in this study, we used our stochastic three-dimensional coarse-grained mesoscale model, that has been proven capable of accurately reproducing experimental time-course saccharification data (29), and we implemented a detailed Sobol’s sensitivity analysis. For this purpose, we created a pipeline combining our stochastic biophysical model together with the Python library SALib (37, 38). Specifically, we systematically evaluated the influence of our model’s parameters on the process of saccharification by comparing simulated time-courses to experimental ones, for two sets of data from the literature (30, 31). Both of them were collected for the enzymatic saccharification of corn stover biomass following three different pre-treatment intensities. For each of the six conditions analysed, we identified the same five out of nineteen model input parameters, which have the largest impact on the saccharification dynamics. These parameters were classified into two categories: those relating to the activity of cellobiohydrolase, and those associated with the crystallinity of cellulose. The consistency of our results across different pre-treatment conditions supported the hypothesis that the same five parameters may play major roles in determining saccharification dynamics in other conditions. We confirmed this assumption for each of the key parameters identified computationally by deriving analytical calculations on a purely cellulosic substrate that accounts for crystallinity and enzyme inhibition by end products. With this mathematical approach, we also elucidated the ranges of the parameter values for which it is expected to find the same key parameters like the ones we identified here. Thereby, we specified the extent of the parameter space in which our results can be generalised, in particular concerning the enzyme kinetic rates of each of the cellulases. We find that for the six conditions considered, the digestion by CBH is the time-limiting step, which is interesting to note since, typically, very little information is available on the composition and kinetics of the commercially available enzyme cocktails used to perform saccharification experiments. This result might even be useful for improving such cocktails, for instance in terms of enzyme variants.

The two pre-treatment procedures applied to the experimental datasets considered here differently modify the biomass, which leads to significant differences in how the saccharification dynamics is influenced by the crystallinity fraction of cellulose. By analysing the profiles of the saccharification curves for the best fits of the experimental data and in case of slight deviations in the value of the cellulose crystallinity fraction, we also pinpointed which part of the time-course is impacted by crystallinity, and confirmed that the digestion dynamics is clearly split into two consecutive phases, with the amorphous bonds being digested first, followed by the crystalline ones. Consistently for both experimental datasets, we showed that for the best fits, the Sobol’s index of the cellulose crystallinity fraction increases with the glucose conversion at 72 hours, while the value of that parameter is instead negatively correlated with the glucose conversion at 72 hours. These two findings both stress the importance of reducing the crystallinity of the substrate in order to increase the final glucose conversion. They also highlight that pre-treatment methods solely focussed on disrupting the crystalline order of the cellulose polymer chains can significantly increase the glucose conversion yield, which may ultimately contribute to advising the design of future pre-treatment techniques.

Noticeably, the deep sampling of the parameter space performed in this study revealed that, in some conditions, clusters of good fits do no converge towards the actual best fit, whose value of cellulose crystallinity fraction is clearly outlying. This urges modellers to carefully navigate the parameter space since seemingly good fits can return very inaccurate parameter values, even for the most impactful parameter of the system. Finally, the results presented will allow prioritising the most important parameters when searching the parameter space, which will certainly enhance the speed of our fitting algorithm, which is so far computationally expensive because of all the parameters being equivalently considered.

## DATA AND CODE AVAILABILITY

The simulation and analysis codes of the model, together with the scripts for reproducing the figures, are provided in the GitLab repository https://gitlab.com/partho9791/data_used_for_sensitivity_analysis_for_pcwsm.

## AUTHOR CONTRIBUTIONS

**PSD:** wrote the manuscript equally with AR, on the basis of a preliminary draft prepared by JT. He produced figure 1 and derived the mathematical calculations of the analytical approach with the support of AR. **JT:** performed the computational approach, and produced the associated figures. She prepared a preliminary draft of the manuscript. **AR:** contributed to derive the mathematical calculations of the analytical approach, conceptualised the work, provided computational resources, supervised the work as well as the entire manuscript preparation, and obtained the funding for the projects PREDIG and EtransColi that she both coordinates. Equally with PSD, she wrote all the content of the manuscript, on the basis of a preliminary draft prepared by JT.

## ACKNOWLEDGMENTS

The authors thank Dr Holger Klose and Dr Philipp Michael Grande from the Institute of Bio- and Geosciences 2 at the Forschungszentrum Jülich (Germany), for fruitful discussions.

## SUPPLEMENTARY MATERIAL

